# Reading the gut microbiome by its fermentative engine and host-facing channels reveals dysbiosis and defines eubiosis

**DOI:** 10.64898/2026.06.09.731167

**Authors:** Matteo Soverini, Ashkan Lotfollahzadeh, Laura di Rito, Antonella Padella, Barbara Santacroce, Andrea Marcante, Cecilia Monaldi, Alena Velichevskaya, Elisa Viciani, Andrea Castagnetti

**Affiliations:** Wellmicro S.r.l., Via Antonio Canova 30, 40128 Bologna, Italy

## Abstract

Taxonomy gives only a partial description of a gut microbiome, since phylogenetically distinct communities encode much the same metabolism and composition is confounded by lifestyle and geography. Here we introduce TAGMOS, an annotation-verified enzymatic architecture aimed at reading gut-microbiome function, built on a fermentative engine and a set of host-facing channels. The engine reports whether the community disposes of fermentative hydrogen through anaerobic sinks or through respiration, and it is the causal driver of the state: antibiotics drive it to an oxidised, respiratory state while faecal transplantation restores the fermentative milieu, and across 73 cohorts the oxidised state is where facultatively anaerobic *Enterobacteriaceae* bloom. We observe that the engine’s disease signal lies in the tail of a population, and a threshold criterion detects disease that a comparison of means overlooks, whereas the channels describe condition-specific mechanistic signatures. Engine and channels transfer across pipelines and populations where taxon-based indices fail. Combining engine and specific channels defines a portable, quantitative measure of eubiosis and dysbiosis, and shows that this state is causally downstream of the community’s metabolic structure.

## Introduction

The gut harbours a microbial community whose influence on human physiology is now recognised across metabolic, immune, hepatic, oncological and neurological disease^1,2^. Its state is usually assessed through taxonomy, to detect which organisms are present and in what proportions - an approach that has well mapped these communities but says little about what they can do. Phylogenetically distinct ecosystems encode and carry out much the same metabolism, so that individuals who differ at the taxonomic level can still share a functional core^3-5^. Taxonomic composition is, moreover, shaped strongly by lifestyle and geography^6^, so the same community structure can have a very different physiological meaning in different populations and individuals. Seen at this layer, the ecosystem is therefore both variable between individuals and disconnected from the metabolic function through which it acts on its host - the dimension through which it could be understood in more detail and, hypothetically, made actionable.

In the present work we interpret the microbiome through the conserved functions it encodes in its metagenome rather than through the taxonomic composition, and reconstruct this functional layer into TAGMOS, an annotation-verified architecture of two coupled parts: a fermentative engine, which summarises how the community disposes of the hydrogen it produces (through anaerobic sinks or through respiration), and a set of channels, which report the host-facing products of its metabolism. Each enzyme enters the framework when specific criteria are met: it must be verified against its curated reaction, mark a rate-limiting or committed step of that reaction, be evolutionarily conserved, and be reliably detectable across bioinformatics pipelines.

The TAGMOS engine captures the redox economy of the colon, the thermodynamic balance on which fermentation relies. In the healthy, hypoxic gut, the *β*-oxidation of butyrate by colonocytes consumes epithelial oxygen and maintains the anaerobiosis that promotes fermentation by obligate anaerobes. When the environment raises epithelial oxygenation and nitrate availability, respiration becomes the more profitable strategy and facultatively anaerobic *Enterobacteriaceae* expand^7-10^. On this view, a disturbed community is not disordered but has adapted to an alternative, stable, respiratory energy state imposed by its host. Interpreted in these terms, we show, the community proves more reproducible and clinically interpretable than when using the underlying taxonomy. The engine responds to causal manipulation and marks the respiratory, oxidised state in which those facultative anaerobes bloom, while the channels resolve the mechanism of individual conditions. Ultimately, we show that eubiosis is mostly a thermodynamic consequence of this redox balance: a portable and quantitative measure of it can be composed from the engine and specific channels, and it proves causally downstream of how the community runs its metabolism.

## Results

### An annotation-verified functional architecture

TAGMOS represents the conserved functional layer of the gut consortium as a defined set of axes, each computed from enzyme functions that are verified against their curated reaction and selected for being rate-limiting or committed steps, for evolutionary conservation, and for detectability across profiling pipelines (Fig. 1A; Methods). Its engine is the redox economy of the community: the balance between the fermentative sinks of microbial hydrogen (acetogenesis, hydrogenotrophic methanogenesis and sulfidogenesis) and the respiratory alternatives that displace them when the ecosystem shifts towards aerobic respiration and nitrate reduction. Alongside this redox balance sit the community’s saccharolytic capacity, the degradation of dietary fibre, and the flavin-based redox cofactor that powers anaerobic fermentation. Finally, the channels report the diffusible products through which the community acts on its host.

**Figure 1.**
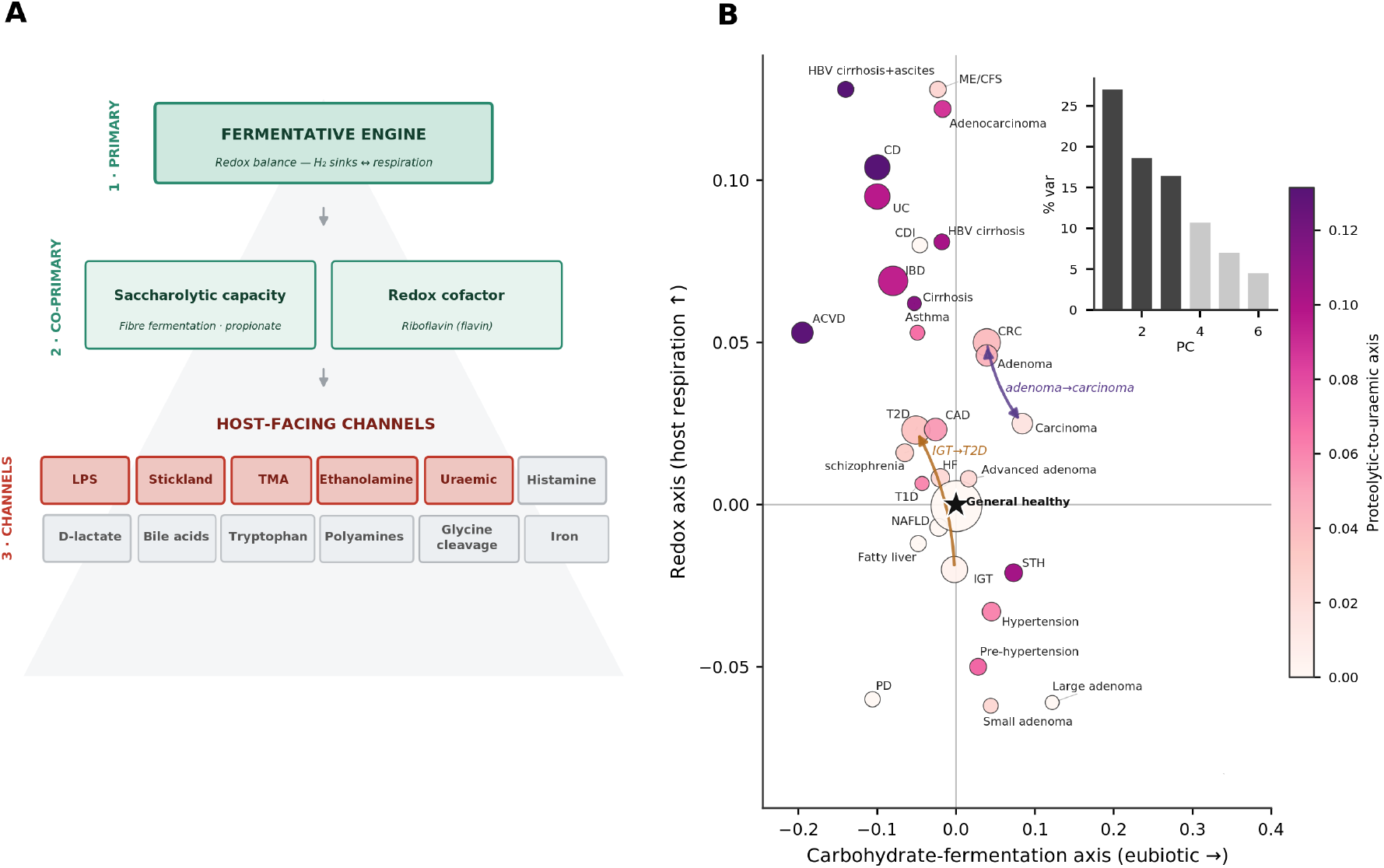
An annotation-verified functional architecture.. **A**, The TAGMOS architecture as a hierarchy: a fermentative engine at the apex - the redox balance of hydrogen disposal, with the saccharolytic and redox-cofactor co-primary axes - widening to the host-facing channels at the base, and the butyrate- and mucin-producing taxonomic guilds corroborating the functional read (not shown). Each axis carries a valence: the engine and its co-primary axes are protective (green), the danger channels that enter the index negatively are red, and the remaining context-dependent channels are grey. A single taxonomy-free composite, EUBIO, weighs the protective engine against the danger channels. **B**, The transdiagnostic metabolic landscape: each condition of cMD positioned on the carbohydrate-fermentation axis (eubiotic to the right) and the redox axis (host respiration upward), colour giving the third, proteolytic-to-uraemic axis and marker size the number of cases; arrows trace the graded colorectal (adenoma→carcinoma) and metabolic (impaired tolerance→type 2 diabetes) sequences. Inset, the variance explained by successive principal components, three axes carrying 62%. Abbreviations (B): ACVD, atherosclerotic cardiovascular disease; BD, Behçet’s disease; CAD, coronary artery disease; CD, Crohn’s disease; CDI, *Clostridioides difficile* infection; CRC, colorectal cancer; HBV, hepatitis B virus; HF, heart failure; IBD, inflammatory bowel disease; IGT, impaired glucose tolerance; ME/CFS, myalgic encephalomyelitis/chronic fatigue syndrome; NAFLD, non-alcoholic fatty liver disease; PD, Parkinson’s disease; STH, soil-transmitted helminthiasis; T1D, type 1 diabetes; T2D, type 2 diabetes; UC, ulcerative colitis.

The resulting architecture is compact: across the conditions of curatedMetagenomicData (cMD)^11^, the adjusted functional signatures resolve onto only three near-orthogonal metabolic axes (Fig. 1B): a redox axis of host-driven respiration, a carbohydrate-fermentation axis, and a third axis of proteolytic-to-uraemic fermentation, the conversion of protein to toxins, which carries most of its variance independently of the other two. These three axes are not the whole of the TAGMOS architecture but they are the low-dimensional coordinate system onto which the disease space mainly projects, and the signature specific to any single condition is drawn from the full set of channels, as shown below. The metabolic state that disease perturbs is set by a few coupled, rate-limiting reactions, but because the surrounding functions co-vary with them, the same shift is mirrored across the whole functional layer rather than contained specifically in a subset of markers. A small, well-chosen set of enzymes therefore captures it as fully as a large one, and random enzyme sets, inside or outside any curated channel, discriminate disease almost as well as the curated panel (cross-validated area under the curve ≈ 0.60 for random sets against 0.63 for the curated one; Extended Data Fig. 1). The discriminating information is therefore not the property of any exclusive feature list but is available to any sufficiently broad sampling of conserved functions. The TAGMOS architecture is, in our view, the minimal structure that reads this redundant signal into compact, mechanistically interpretable and correctly annotated coordinates (Methods).

### The fermentative engine tracks the state and responds to its causes

The engine reports whether the community is oriented towards fermentation or respiration, and its disturbance in disease is detected most sensitively not from the average of that balance but from whether a sample has crossed into a respiratory, oxidised state - bypassing a threshold rather than shifting away from the mean. The balance is expressed as a within-sample log-ratio of the hydrogen sinks to the respiratory routes, a ratio that does not depend on cohort-specific standardisation and therefore keeps its meaning across populations (Methods; Extended Data Fig. 3), and defines an oxidised niche whenever respiration outweighs the sinks. Seen this way, the engine’s disease signal is a tail phenomenon that a comparison of means fails to capture (Fig. 2A): in inflammatory bowel disease (IBD) the continuous ratio is unmoved (odds ratio 0.80 per standard deviation, P = 0.14), yet the fraction of subjects in the oxidised niche separates cases from controls (10.6% of cases against 3.9% of controls, gate odds ratio 2.0, P < 10^−4^), and in cirrhosis nearly half of cases occupy the oxidised niche against a sixth of controls. Colorectal cancer (CRC), by contrast, is engine-neutral on both readings, its disturbance being carried instead by the channels, so that the engine and the channels report on different diseases through different parts of the same architecture.

**Figure 2.**
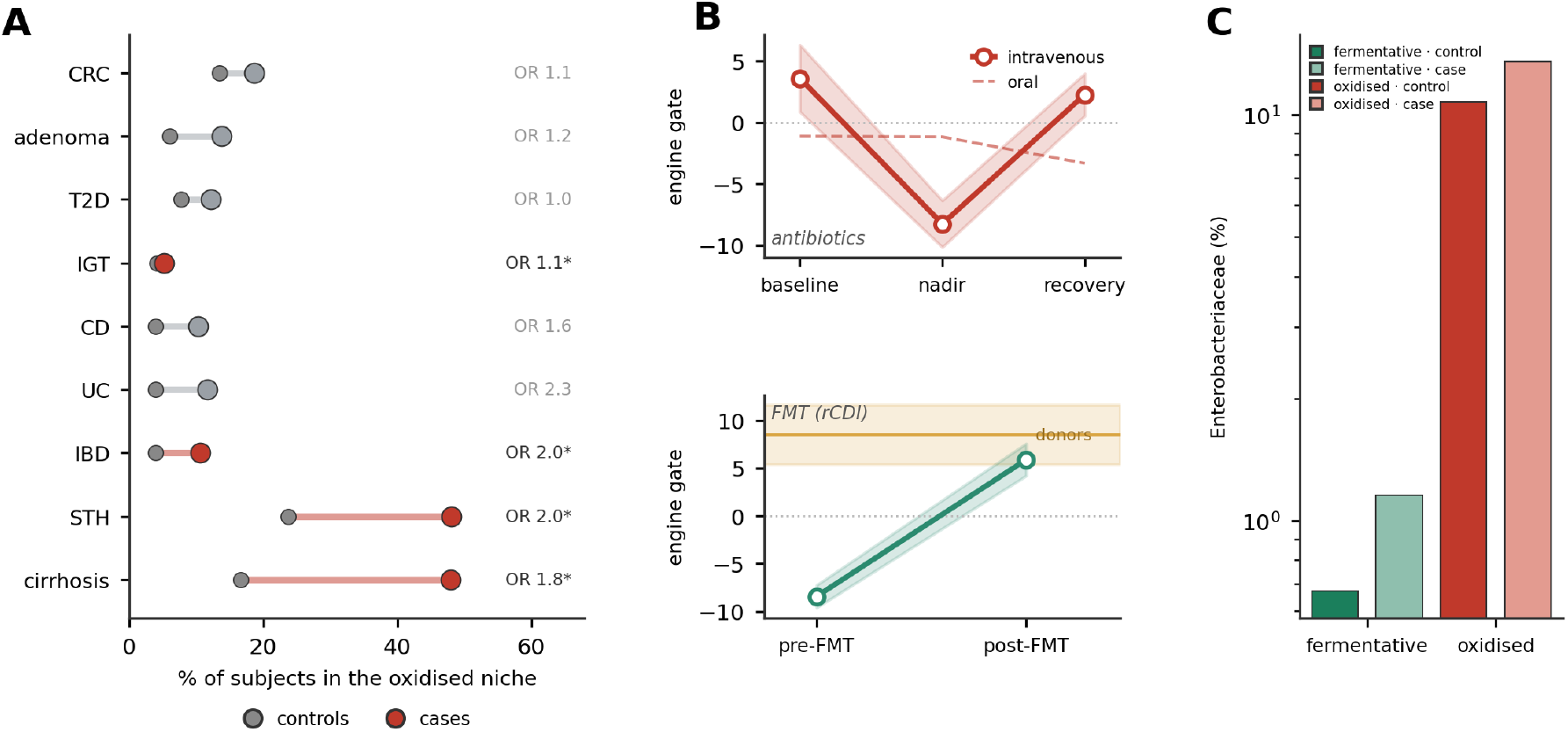
The fermentative engine tracks the state and responds to its causes.. **A**, The engine dysbiosis as a threshold phenomenon: the fraction of subjects in the oxidised niche (respiration outweighing the fermentative sinks) separates cases from controls even where the mean sink-to-respiration ratio does not, with the gate odds ratio annotated (asterisk, gate P < 0.05). **B**, The engine gate (within-sample log-ratio of sinks to respiration) under perturbation, shown as the mean trajectory with its standard error: antibiotics drive it down, steeply for an intravenous course and only shallowly for an oral one (top), while faecal microbiota transplantation for recurrent *Clostridioides difficile* infection lifts it from a pre-treatment low towards the donor level (band, bottom). **C**, *Enterobacteriaceae* relative abundance across the four niche- by-group strata (log scale): the fermentative niche in green and the oxidised niche in red, each split into controls (dark bars) and cases (light bars). These facultative anaerobes are roughly ten- to twenty-fold more abundant in the oxidised than in the fermentative niche, and their case-versus-control excess is concentrated in the oxidised stratum; the engine’s oxidation tracks their abundance across studies (partial correlation -0.41, negative in 72 of 73).

We observe that this threshold behaviour is not peculiar to the engine but a general property of the architecture. Applied across the channels and conditions of cMD, roughly 25% of the detectable disease associations appear only as a threshold and are missed altogether by a comparison of means, while about 70% carry a threshold signal of some strength, and these threshold-only associations reproduce across studies at least as well as the mean-based ones (sign concordance 0.97 against 0.94; Methods; Extended Data Fig. 4). A substantial part of the disease signal in the gut microbiome is therefore a tail phenomenon - a subgroup crossing a threshold rather than a whole population shifting on average - which may help to explain why associations sought as differences in the mean have often proved difficult to replicate.

Because the engine responds to the interventions that are well known to perturb gut ecology, it can be taken as a cause of the state rather than a correlate of it (Fig. 2B). Antibiotics drove the community into the oxidised, respiratory state, and the depth of the collapse tracked the severity of the exposure: within subjects, an intravenous course collapsed the engine from a fermentative baseline to a deeply oxidised nadir, at which seven in ten samples had crossed into the oxidised niche, before it recovered once the course ended^12^, whereas a milder oral course produced only a shallow, transient decline^13^. Faecal microbiota transplantation for recurrent *Clostridioides difficile* infection did the reverse, lifting the engine from a pre-treatment state in which most samples were oxidised back towards the fermentative level of the donors^14-16^. Indications in which transplantation does not reliably rebuild the community, such as metabolic syndrome and IBD^17,18^, left the engine unmoved.

A reverse validation of the oxidised niche can be drawn from the organisms that populate it (Fig. 2C): across seventy-three studies the engine’s oxidation tracks the abundance of *Enterobacteriaceae*, the facultative anaerobes that expand by respiring host-derived electron acceptors. As the fermentative log-ratio falls, these organisms rise (partial correlation -0.41 overall, negative in 72 of 73 studies). They are roughly ten-to twenty-fold more abundant in samples lying in the oxidised niche than in fermentative ones, and their excess in cases over controls is concentrated in that oxidised stratum: their relative abundance from taxonomic profiling reaches 13.6% in cases against 10.7% in controls in the oxidised niche, but only 1.2% against 0.7% in the fermentative one. At the level of conserved function, then, the engine captures the same oxygen-driven transition that taxonomic and mechanistic studies have described one organism at a time^7-10^.

### The channels qualify disease by mechanism

The third and more granular level of the TAGMOS architecture is its host-facing channels, through which the mechanism of a disease can be traced back to the specific functions it has perturbed. Protective channels are consistently higher in fermentation-oriented communities and fall as the community is disrupted and shifts towards respiration. Damage channels are the mirror image, consistently higher in oxidative communities - the proteolytic-to-uraemic group of Stickland proteolysis, ethanolamine utilisation, p-cresol and trimethylamine production and endotoxin biosynthesis, which rise together across disease. Context channels take their sign only once the condition is known, rising in some diseases while falling in others: hydrogen sulfide, secondary bile acids, histamine and the polyamines are protective up to specific definable levels but damaging in excess, qualifying a disease rather than grading its severity (Fig. 3B, right-hand valence strip). Across inflammatory and hepatic disease, with every effect computed within its own study and adjusted for age, sex, body mass, antibiotics, sequencing depth and functional richness (Methods), a single proteolytic-to-uraemic axis - the fermentation of protein to toxins^19^ - rose consistently and ordered the conditions from IBD through to cirrhosis, carrying its proteolytic, ethanolamine, uraemic and trimethylamine channels with it (Fig. 3A,B). The signal was a gradient rather than a total switch, and it tracked both the colorectal adenoma-to-carcinoma sequence and the progression from normal glucose tolerance to type 2 diabetes (T2D). Along the colorectal sequence the propionate, tryptophan and trimethylamine channels rose with disease stage (Spearman *ρ* = +0.30, +0.27 and +0.26, respectively), and along the diabetic progression the uraemic, ethanolamine and endotoxin channels rose together (*ρ* = +0.19, +0.18 and +0.17, respectively). On this basis, a disease is better described as a multi-step trajectory than as a single case-control contrast.

**Figure 3.**
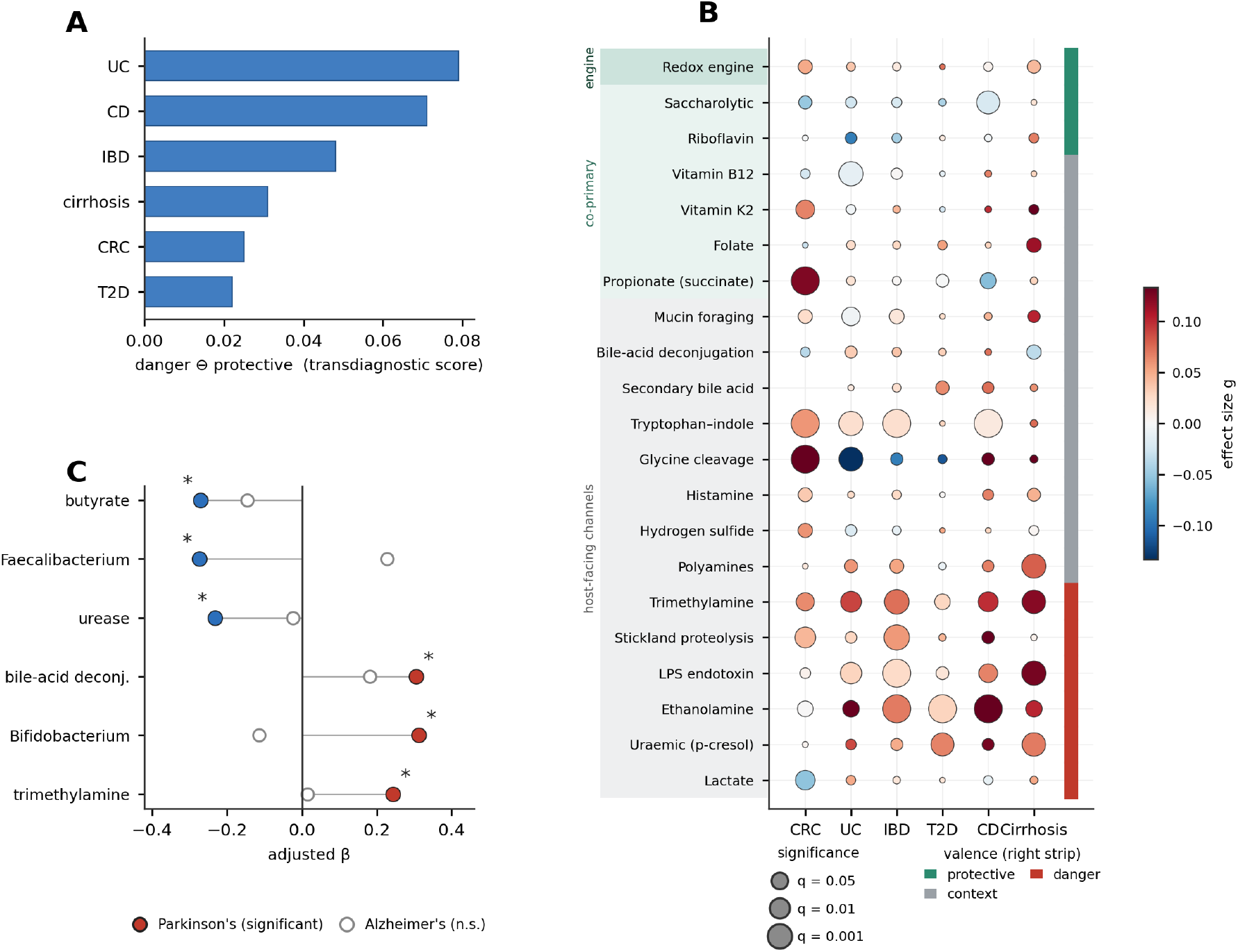
The channels qualify disease by mechanism.. **A**, Transdiagnostic ordering of conditions on the proteolytic- to-uraemic axis (danger minus protective). **B**, Channel-by-condition bubble matrix across six conditions of cMD: colour gives the effect size (Hedges’ *g*, red = higher in cases), bubble area the significance (-log_10_ *q*) and the right-hand strip the channel valence (protective, context or danger); rows follow the architecture hierarchy (protective engine, then context, then damage) and conditions are clustered by similarity. Axes excluded from the framework for unreliable direction or poor portability (GABA, tyramine, NAD) are not shown; iron acquisition, a qualitative presence flag without a continuous score, is likewise omitted here and appears only in the architecture (Fig. 1A). **C**, Neurodegeneration: covariate-adjusted channel effects for PD (filled) and AD (open); PD is functionally dysbiotic and survives adjustment for constipation, whereas AD is not.

That CRC trajectory carried the strongest of all the single-disease signatures. The microbial glycine-cleavage system - which feeds glycine into one-carbon folate metabolism^20^ - was elevated in CRC across all 11 cMD studies (gate odds ratio 2.83), was robust to covariate adjustment, and rose in step from adenoma to carcinoma (gate odds ratio 2.07 and 2.83, respectively; adenoma P = 5.3 × 10^−4^, carcinoma P = 2.3 × 10^−6^). The signature was specific to colorectal neoplasia, being null or even reversed in the other conditions, and it was independent of the fibrolytic and trimethylamine axes, holding its association after adjustment for the butyrate taxonomic guild and trimethylamine (odds ratio 1.61, P = 6 × 10^−4^). This specificity fits the metabolic direction it marks, the serine-and-glycine one-carbon metabolism known to supply nucleotides to proliferating tumours^20^. As with the other single-disease signatures, we report it as a reproducible association, not claiming it as a cause.

When applied to neurodegeneration, the channels separate two diseases while withstanding the confound most likely to mislead them. Parkinson’s disease (PD) is reported as dysbiotic by taxonomic surveys, and animal transfer of a PD microbiota worsens motor deficits and α-synuclein pathology^21,22^, yet the obvious confound is that the disease slows intestinal transit. Computed on the public cohort of Wallen and colleagues^23^ and adjusted again for constipation, age, sex, body mass, antibiotics, sequencing depth and functional richness, the phenotype survived in both its functional and its taxonomic components: butyrogenic capacity, together with Faecalibacterium, fell, while bile-acid deconjugation and Bifidobacterium rose (Fig. 3C). Alzheimer’s disease (AD) provided the negative control: in a cohort spanning normal cognition to dementia^24^, the measure neither separated cases from controls nor followed the clinical progression.

### Eubiosis is a consequence of the engine and specific channels

Having characterised how the engine tracks the state and how the channels resolve disease, we can define a single measure of eubiosis on this architecture and ask where it stands in the causal order. We therefore introduce EUBIO, a taxonomy-free scalar that weighs the protective engine against the channels that carry host damage, calibrated once on a reference of healthy adults and computed on the conserved functional layer rather than fitted to case-versus-control data (Methods). Its response to perturbation places it downstream: it moves as the engine does, collapsing under antibiotics and recovering after transplantation in proportion to the effectiveness of the intervention, and when its within-subject change is decomposed into an engine contribution and a channel contribution, the engine accounts for about two thirds of it (0.63 of the paired change, and 0.73 under the intravenous antibiotic; Fig. 4A,B). Eubiosis is therefore a downstream, derived measure of the community’s metabolism within the TAGMOS architecture, and its causal movement is carried for the most part by the engine.

**Figure 4.**
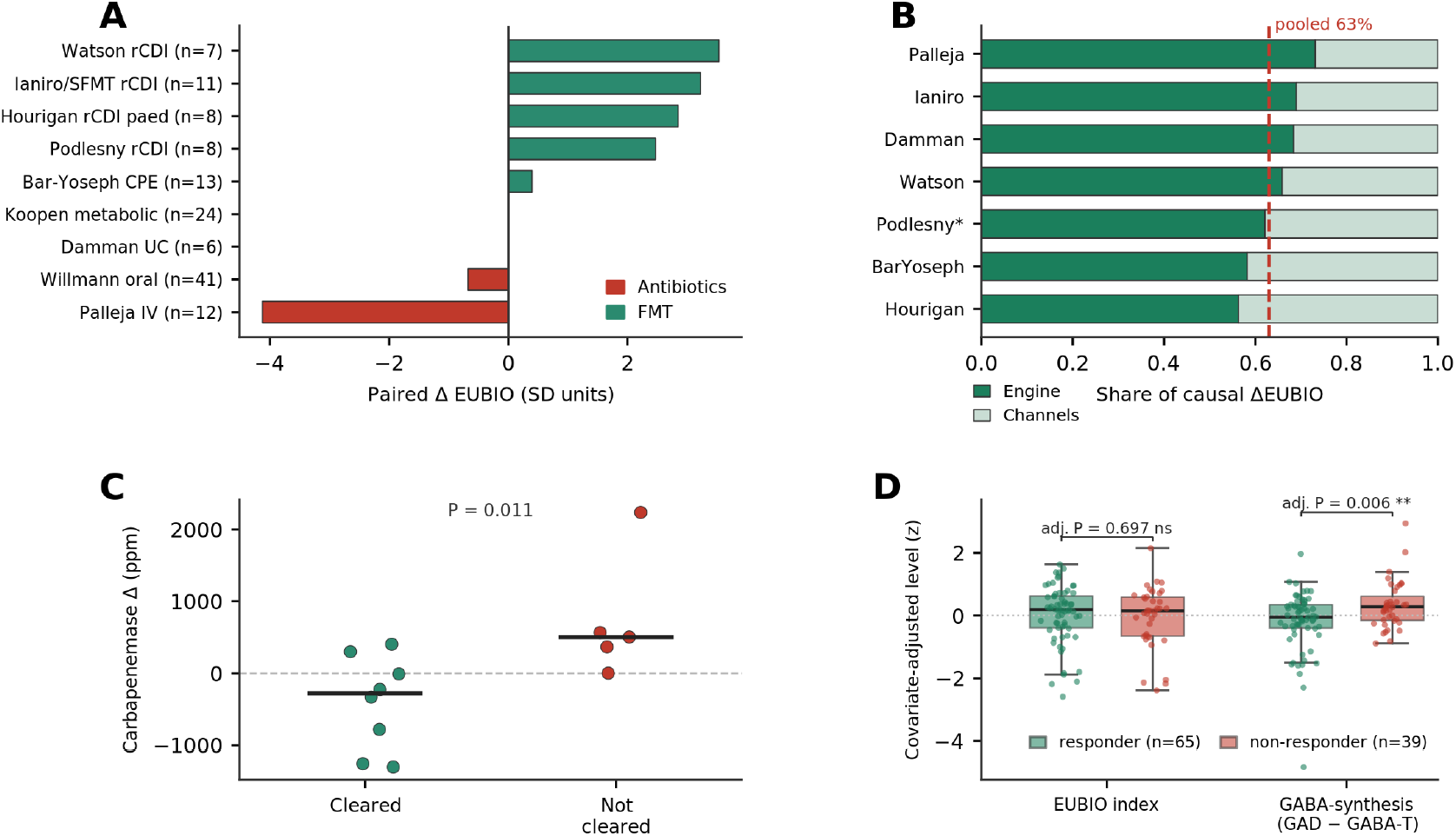
Eubiosis is a consequence of the engine and its channels.. **A**, EUBIO responds to causal manipulation, falling under antibiotics and rising after transplantation by an amount set by the indication. **B**, Decomposing the within-subject change in EUBIO into an engine and a channel contribution: the engine carries about two thirds of it across cohorts (dashed line, pooled 63%). **C**, Clinical meaning: in decolonisation, the carbapenemase enzyme (EC 3.5.2.6) falls in patients who cleared the organism and rises in those who did not (*P* = 0.011). **D**, Specificity and function-resolution in melanoma checkpoint immunotherapy (PRIMM baseline cohort, *n* = 104). The global EUBIO index does not separate responders from non-responders (covariate-adjusted *P* = 0.70), whereas the microbial GABA-synthesis channel (glutamate decarboxylase minus GABA-transaminase) is lower in responders (covariate-adjusted *P* = 0.006). Boxes show covariate-adjusted levels (responders green, non-responders red; adjusted for age, sex, BMI, antibiotics and depth). The direction recurs across independent melanoma cohorts (three of three on durable benefit) and is coherent with microbial GABA restraining anti-tumour immunity - a hypothesis-grade signal, not a biomarker.

Beyond its causal standing, the measure is clinically relevant. In a cohort undergoing faecal transplantation to decolonise carbapenemase-producing *Enterobacteriaceae*^25^, it tracked the recorded clinical outcome, rising in patients who cleared the organism (a paired gain of +1.29 standard-deviation units in 8 of 8 patients), while the carbapenemase enzyme itself (EC 3.5.2.6) fell in those patients and rose in those who did not (P = 0.011), a more direct genomic signature of eradication (Fig. 4C). EUBIO is also specific to effective eubiosis and does not simply reward any microbiome that happens to help the host by a route other than gut ecology: in melanoma patients starting checkpoint immunotherapy, whose benefit is immunemediated rather than ecological^26-28^, the global index does not separate responders from non-responders (covariate-adjusted P = 0.70; Fig. 4D), whereas resolving the same community by function isolates a specific and reproducible baseline signal, which we develop below. In coverage-adequate repeated samples the scalar was a stable individual characteristic (intraclass correlation 0.47; PRJNA664754).

### The architecture is portable across pipelines and populations

Portability requires that TAGMOS, and the EUBIO scalar built on it, remain stable across a change of profiling pipeline and a change of population, and we tested both scenarios. On paired samples processed by two independent pipelines, our assembly-based Wellmicro pipeline (WMP; Methods) and the HUMAnN3 pipeline of bioBakery 3^29^, the fermentative core is detected in every sample by both pipelines, the engine’s ranking transfers between them, and freezing the calibration on the reference does not alter that ranking (Spearman 1.00 for the core, 0.88 for the full index; Extended Data Fig. 3). The sparser single-enzyme host-facing channels are detected only where sequencing coverage allows. On short-read pipelines their signal is too sparse to be interpreted on its own and is used only to support the interpretation of the densely detected core. Carried across populations (Fig. 5A), the architecture-based EUBIO scores traditional-diet communities as eubiotic (Hadza +1.56; Andean and Amazonian traditional populations +1.51, on samples of usable depth)^30,31^, where the taxonomic components of conventional indices, defined on Western clinical or beneficial taxa, score them as deeply dysbiotic. Within a single population, EUBIO separates false dysbiosis from real, declining monotonically across the Sardinian lifespan^32^ from young adults (+1.11) through the elderly (+0.56) to centenarians (-0.92), tracking the dysbiosis of extreme old age.

**Figure 5.**
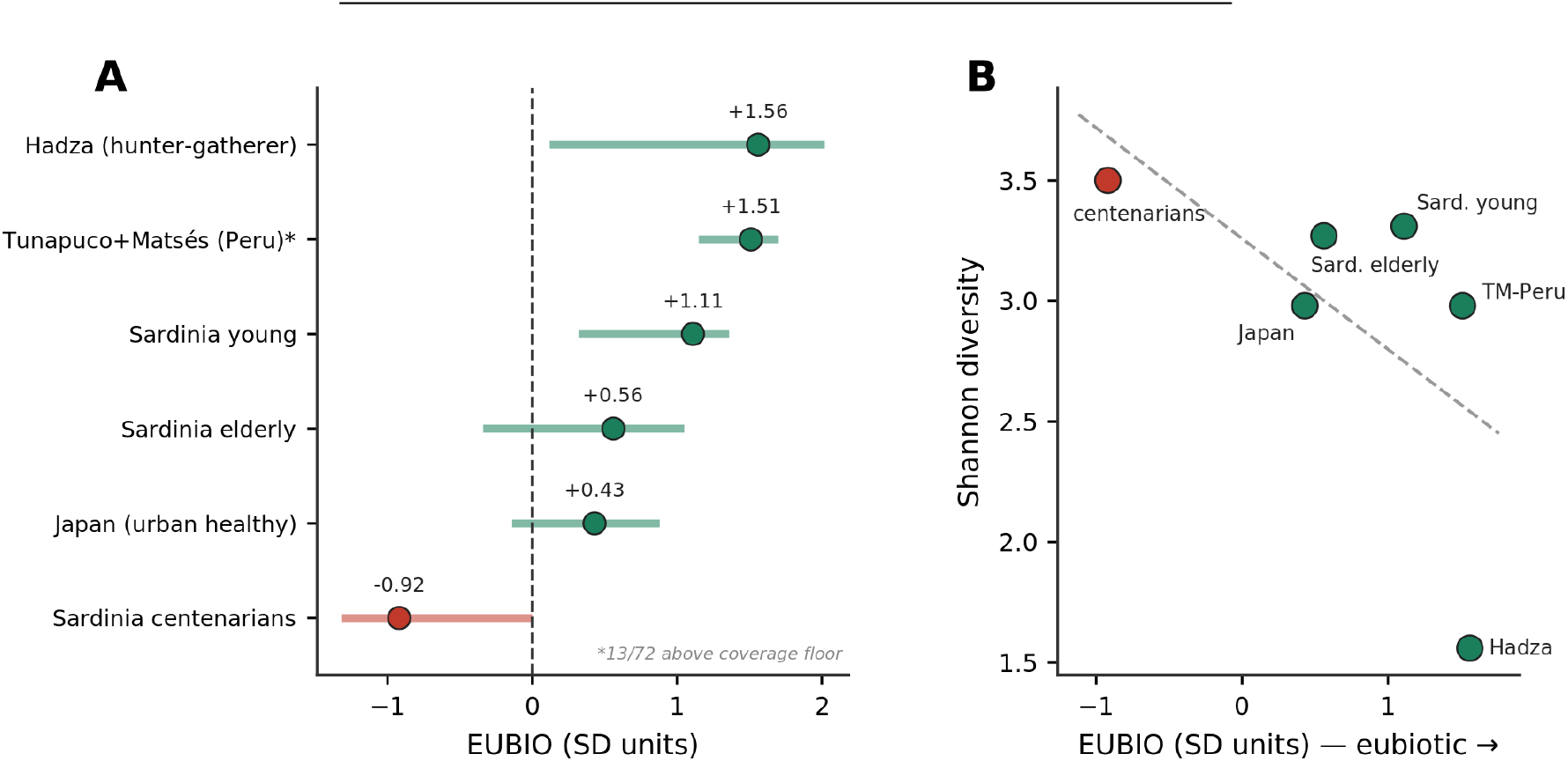
The architecture is portable where taxonomy-based metrics fail.. **A**, EUBIO by population (median, interquartile range); traditional-diet and healthy populations score eubiotic while the Sardinian lifespan declines to a genuine dysbiosis in centenarians. **B**, Shannon diversity plotted against EUBIO across the same populations: diversity is highest in the most dysbiotic group and lowest in the most eubiotic, inverting the functional ranking. —

### Taxonomy-based metrics fail where function holds true

Eubiosis and dysbiosis have long been estimated through taxonomic proxies - *α*-diversity and indices built on panels of health-associated genera - on the assumption that a richer or more “normal” community is a healthier one. Across the populations examined above, however, these proxies sometimes point the wrong way: Shannon diversity is highest in the Sardinian centenarians, the most dysbiotic group, and lowest in the Hadza, the most eubiotic, the exact inverse of the functional ranking, because a diverse community is not for that reason a healthy one (Fig. 5B). An index built on a fixed panel of health-associated genera is no better, scoring the hunter-gatherers lowest of all because their communities lack the Western genera it is built to count (Fig. 5B). Diversity and this functional state are complementary but not interchangeable, and the divergence between the two holds not only for a state but for a treatment response, where taxonomy again fails and function does not. In a cohort treated with immune-checkpoint inhibitors, in which taxonomic associations with response have themselves proved largely cohort-specific and non-transferable^28^, EUBIO did not separate responders from non-responders at baseline. A single functional channel, however, did. Microbial GABA synthesis, quantified from glutamate decarboxylase (EC 4.1.1.15), was lower in responders (adjusted P = 0.006), reproduced within each of three independent melanoma cohorts on a different pipeline^11^, and was independent of taxonomy: no species carried a reproducible responder signature, the GABA-producing genera trended the opposite way, and the enzymatic association was undiminished after adjustment for their abundance (coefficient from -0.44 to -0.48). Microbial GABA synthesis is an immunosuppressive input to antitumour immunity, acting through interleukin-10-secreting macrophages that restrain cytotoxic T cells^33^. The association is modest and reported as a mechanistic hypothesis, but it shows that the transferable signal is carried by conserved function and not by the taxa that encode it.

## Discussion

Taxonomy has mapped the gut community in extraordinary detail, yet it remains a partial guide to what that community does, because taxonomically divergent communities converge on the same metabolism and because composition is confounded by lifestyle and geography^3,5^. Interpreted instead through the conserved functional layer, and organised into a fermentative engine and a set of mechanistic channels, the community becomes measurable, comparable across populations and mechanistically interpretable, and the long-standing difficulty of defining a dysbiotic state^34-36^ dissolves into a property that can be measured.

That the engine is the right primary object of measurement rests on a chain that closes causally. Antibiotics drive the community into the oxidised, respiratory state and transplantation restores the fermentative condition, placing eubiosis and dysbiosis as two energetic states of the host-microbe coupling rather than as order and its collapse. The oxidised state is the respiratory niche in which facultative anaerobes bloom, exactly the oxygen biology described one organism at a time. And when the eubiosis scalar moves under perturbation, most of that movement is driven by the engine. Treating the engine as a threshold rather than a mean has its relevance, because the transition is a tail phenomenon: a minority of a diseased population crosses into the oxidised niche, and averaging over the majority that has not conceals it. This proved to be a general property of the architecture rather than a peculiarity of one axis. Across cMD, about a quarter of detectable disease associations are threshold phenomena invisible to a comparison of means, and they replicate across studies at least as well as mean associations do. Much of the microbiome field’s difficulty in reproducing disease associations may therefore be methodological rather than biological - a matter of testing for a shift in the mean where the signal lies in the tail - and profiling the community through population-invariant thresholds recovers a class of association that mean-based analysis discards.

Reading the microbiome as function rather than membership changes what a clinical result can be. Because the engine and its channels are mechanistic and calibrated on health, a disturbed community points to the specific reaction that has moved - the respiratory shift behind an *Enterobacteriaceae* bloom, one-carbon metabolism in colorectal neoplasia, or microbial GABA synthesis before checkpoint immunotherapy - rather than to a list of taxa whose meaning varies between people. This turns a descriptive survey into a set of candidate mechanisms and targets, and makes the same measurement comparable across populations that share almost no taxa, from industrialised cohorts to hunter-gatherers. It also reframes a persistent difficulty of the field: the poor reproducibility of taxonomic disease associations may reflect, in part, a search for shifts in the mean where the signal is a minority of subjects crossing a threshold, a pattern that a population-invariant, function-level readout recovers.

What makes this measurable is the biological centrality of the processes involved, not the length of the feature list. Hydrogen disposal and the fate of carbohydrate fermentation are among the most conserved transactions in microbiology^37^, and because the state they set is redundantly encoded across the community, it can be captured reliably through a compact set of correctly annotated coordinates rather than an exhaustive panel. Getting that annotation right is what keeps the coordinates faithful, and the same discipline sharpened the disease biology rather than changing its substance. This sets the architecture apart from schemes built to summarise community structure, such as enterotypes^38^ and enterosignatures^39^, and from data-driven health indices that are correlative classifiers of taxa trained to separate cases from controls^40^: here the object is a mechanism, calibrated on health rather than fitted to disease, from which the channels resolve condition-specific signatures and the scalar follows.

These claims are bounded. Only the colorectal signature, across eleven studies, and in part the adenoma sequence, are confirmatory; the remaining single-cohort signatures are mechanistic hypotheses for trial, not biomarkers. The fermentative core transfers across pipelines and populations, but the sparser single-enzyme channels are under-detected on short-read data and are used there only in support; and because the absolute calibration is fixed to a proprietary reference, every result reported here rests on public data through within-subject change, rank or within-study contrast and the population-invariant ratio form of the engine. Within these bounds, TAGMOS offers a mechanistic, portable and causally grounded reading of the gut microbiome, of which a measurable eubiosis is the consequence.

## Methods

### Cohorts and processing

TAGMOS was developed and evaluated entirely on public metagenomic cohorts spanning causal interventions, disease, neurodegeneration and diverse human populations; the proprietary healthy reference enters only as the frozen calibration of the scalar (below). The causal anchors comprise two antibiotic cohorts^12,13^ and cohorts of faecal microbiota transplantation undertaken for eubiosis restoration or decolonisation, spanning recurrent *Clostridioides difficile* infection^14-16,41,42^, decolonisation of carbapenemase-producing and multidrug-resistant organisms^25,43^, metabolic syndrome^17^ and IBD^18,44^. Two cohorts of transplantation from immunotherapy responders served as the test of specificity^27,45^, and one public longitudinal cohort of healthy adults (a six-month test-retest of 20 healthy volunteers; PRJNA664754) as the test of reliability. Transdiagnostic disease biology and the engine-versus-taxonomy analysis used cMD^11^ profiled with bioBakery 3^29^. The neurodegeneration analyses used the public PD cohort of Wallen et al.^23^ together with an AD cohort^24^ reprocessed through our pipeline. The cross-population analysis used Japanese^46^, Sardinian^32^, Hadza^30^, and Andean and Amazonian traditional^31^ metagenomes.

All metagenomes generated or reprocessed in house were passed through the Wellmicro Metagenomics Pipeline (WMP). After adapter trimming and quality filtering with fastp^47^, reads were classified with Kraken2^48^ against a proprietary cross-kingdom database of complete NCBI reference genomes, so that host and other non-bacterial reads are competitively removed in the same step that assigns taxonomy. Function was quantified at the level of Enzyme Commission (EC) numbers by assembling contigs with MEGAHIT^49^, detecting open reading frames with FragGeneScan^50^, and aligning the predicted proteins with DIAMOND^51^ against a curated bacterial protein database at 70% amino-acid identity; every in-house sample was subsampled to a uniform 1.3 Gbp before functional quantification to remove sequencing depth as a source of variation. curatedMetagenomicData is distributed as HUMAnN3 UniRef90 gene-family abundances (in reads per kilobase, RPK) rather than as ready-made EC profiles, and was not reassembled. We converted it to EC coordinates with a mapping built by inverting HUMAnN’s official level-4 EC-to-UniRef90 table (`map_level4ec_uniref90`) into a UniRef90-to-EC dictionary: each UniRef90 family contributed its abundance to every EC it represents (multi-functional families were kept as such rather than forced to a single assignment), abundances were summed over the taxonomic stratification, and the per-study matrices were merged on the union of ECs (5,796 ECs across 22,409 samples). Because HUMAnN3 RPK and the assembly-based WMP counts lie on different scales, each EC was quantile-mapped onto a fixed healthy reference before the scalar was computed; being a monotone per-EC transform, this reconciles scale without altering within-cohort rankings (Extended Data Fig. 3). Where a reaction can be labelled by more than one EC identifier - for example a current and a superseded number - the identifier actually detected in a given pipeline was used, so that the same reaction, and not merely the same code, is compared across pipelines.

### The TAGMOS architecture: engine and channels

TAGMOS is a reconstruction of the conserved functional layer in which every enzyme is mapped to its curated reaction and retained only if it marks a rate-limiting or committed step, is evolutionarily conserved, and is detectable across profiling pipelines; enzymes that are housekeeping, promiscuous or ambiguous for the reaction they are taken to mark are not included, and the full enzyme dictionary is proprietary and is not released. The axes of the architecture were not imposed a priori but recovered from the data: projected across the conditions of cMD, the retained functions collapse onto a small number of near-orthogonal metabolic axes (Fig. 1B), among which the redox engine emerges as the dominant and causally responsive one. Applying the same reaction-level criteria occasionally reassigns an enzyme from the function it is conventionally used to mark to the one it actually catalyses - for instance restricting the endotoxin channel to the committed hexa-acylation of lipid A, or quantifying trimethylamine from choline trimethylamine-lyase (CutC, EC 4.3.99.4)^52^ - with the corresponding reactions described here at the level of chemistry rather than of proprietary enzyme identity.

The **engine** is the redox economy of fermentation. Its core is the balance between the terminal fermentative sinks of microbial hydrogen (acetogenesis, hydrogenotrophic methanogenesis and sulfidogenesis) and the respiratory routes that displace them, aerobic terminal oxidases and nitrate reduction. It is computed continuously and, for a population-invariant measure, as a within-sample log-ratio, log_2_((Σ sinks + 0.1)/(Σ respiration + 0.1)) on abundance ratios, which defines an oxidised niche when respiration outweighs the sinks by a fixed factor. This ratio form does not depend on cohort-specific standardisation and therefore transports across populations. Alongside the redox core, the engine includes the community’s saccharolytic capacity, the degradation of dietary fibre, and a flavin-based redox cofactor. A compact two-enzyme readout of the fermentative-versus-respiratory partition of carbon (EC 4.1.1.49 and EC 2.7.3.9) provides the portable coordinate used for cross-pipeline transfer. The **host-facing channels** capture the diffusible products through which the community’s metabolism reaches the host. Those that carry host damage (lipopolysaccharide endotoxin biosynthesis, Stickland proteolysis, ethanolamine utilisation, uraemic-toxin production and trimethylamine production) are the ones that enter the index negatively, alongside additional context channels (polyamine and iron-acquisition) treated as flags. A small number of channels are taxonomic guilds rather than EC axes, notably the butyrate-producing guild and the mucin-foraging guild, and these are used to qualify disease but are not part of the portable, taxonomy-free core.

### The EUBIO scalar and frozen calibration

EUBIO is a signed, continuous composite that weighs the protective engine against the channels that carry host damage. Its positive pole is eubiotic and its negative pole dysbiotic, and it is expressed in standard-deviation units of the healthy reference. Its per-channel calibration is frozen on a healthy reference defined primarily by clinical self-report on the Wellmicro internal real-world-evidence cohort, and broadened by public healthy anchors with documented clinical status. Because the reference is not public, no reported result depends on it, and this rests on three grounds: the architecture and the sign of each channel are validated on the public causal anchors; every reported effect is a within-subject change, a rank or a within-study contrast on public cohorts; and the ranking is invariant to the frozen calibration, as shown on the public PD cohort, where the fermentative core is perfectly scale-invariant (Spearman 1.00) and the full index nearly so (0.88). Because EUBIO is a signed sum of standardised axis scores, a within-subject change decomposes additively: the engine contribution is the summed change of the fermentative (engine) axes and the channel contribution the summed change of the damage channels, each reported as a fraction of the total absolute change on the paired anchor samples.

### Validation and statistics

For the disease analyses, every comparison was conducted within its own study and never pooled, adjusted for age, sex, body mass, antibiotics, sequencing depth and functional richness (the number of enzymes detected). Concordance was assessed across studies by random-effects meta-analysis and Benjamini-Hochberg correction, and a signature was called confirmatory only when it reproduced across at least three independent studies. The engine gate was evaluated both as a continuous odds ratio per standard deviation of the sink-to-respiration log-ratio and as a threshold, the odds ratio for occupying the oxidised niche, so that tail phenomena invisible to the mean are captured. To test whether such threshold behaviour generalises beyond the engine, every channel-by-condition association was evaluated both as a mean effect (odds ratio per standard deviation) and as a threshold (odds ratio for occupying the dysbiotic tail, defined relative to each study’s own controls), and each was classified as detectable by the mean, by the threshold, or by both; the fraction detectable only through the threshold, and the cross-study sign concordance of each measure, were tabulated across a range of threshold values and false-discovery cutoffs, and were stable to both. The engine’s causal response was measured as the per-phase mean of the log-ratio across the antibiotic and transplantation cohorts, and its mechanistic anchoring as the within-study partial correlation of the log-ratio with *Enterobacteriaceae* relative abundance from taxonomic profiling. The transdiagnostic ordering weighed the damage axes against the protective axes. The disease cascades were the within-study Spearman correlation of each channel with disease stage. The butyrate guild was tested against a negative control of 150 random species panels matched for prevalence. Differences between groups were tested by the Mann-Whitney test, paired changes by the paired *t* or Wilcoxon test, channel associations by covariate-adjusted models with Benjamini-Hochberg correction, and discrimination by the area under the curve, with standardisation within each cross-validation fold.

## Data availability

Every cohort used in this work is public and respective accessions are listed in the Extended Data section. The aggregated cohort- and condition-level TAGMOS outputs (axis effect sizes, gate odds ratios and per-population summaries) are provided as source data with this article. The reconstructed enzyme dictionary and the frozen calibration constants are proprietary and are not released.

## Code availability

The architecture and the definitions used here are described in this manuscript and its Extended Data. Analysis code operating on the public cohorts is available from the corresponding author on reasonable request, and the frozen model is provided as a service through a dedicated portal, currently in development. Until it is online, requests to reproduce the analyses reported here, or to classify specific samples (including blind classification, in which the clinical condition is withheld so that the classifier can be tested independently), can be directed to the corresponding author.

## Acknowledgements

We thank the investigators and participants of all the public cohorts analysed, and the individuals tested through the Wellmicro® service who allowed the research use of their data.

## Author contributions

M.S.: Conceptualization, Methodology, Software, Formal analysis, Investigation, Writing - original draft, Writing - review and editing, Visualization. A.C.: Conceptualization, Methodology, Supervision, Funding acquisition, Writing - original draft, Writing - review and editing. A.L. and L.d.R.: Data curation, Resources. A.V., A.P., B.S., A.M., C.M. and and E.V: Investigation. All authors read and approved the final manuscript.

