## Supplementary material for "Reading the gut microbiome by its fermentative engine and host-facing channels reveals dysbiosis and defines eubiosis": Supllementary figures and tables

#### Extended Data Figures

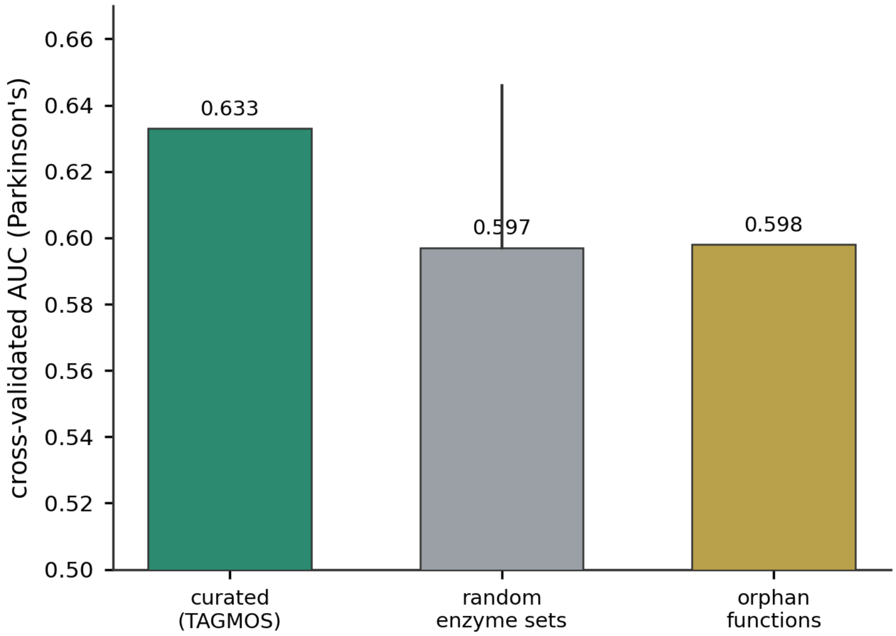


**Extended Data Fig. 1 | The discriminating signal is pervasive..** Cross-validated discrimination of PD by the curated TAGMOS panel, by random enzyme sets of equal size (mean, with the maximum over draws), and by orphan functions outside any curated channel; all discriminate well above chance.

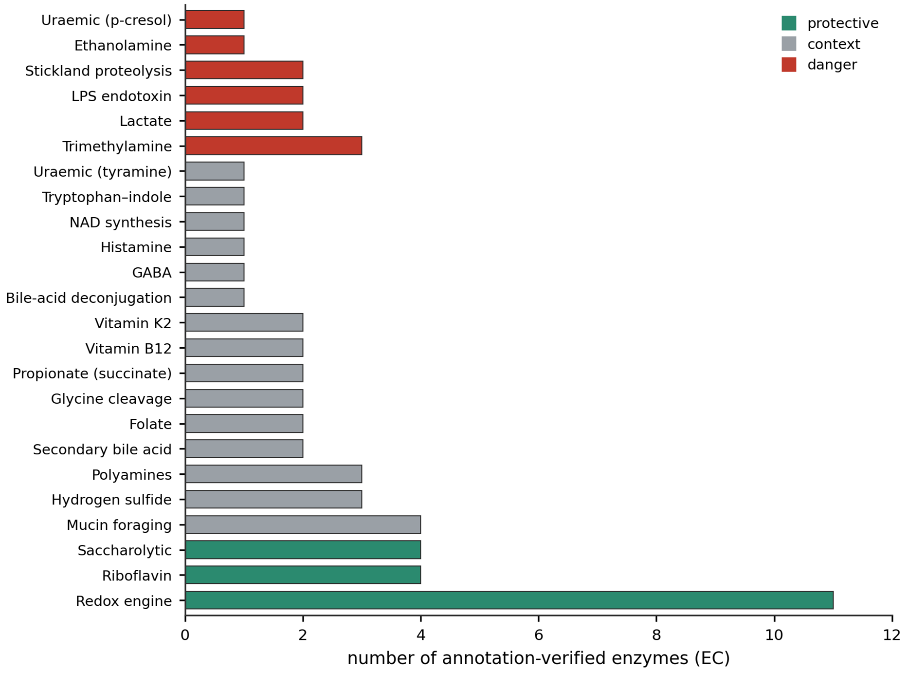


**Extended Data Fig. 2 | TAGMOS architecture: enzymes per axis..** The number of annotation-verified enzymes assigned to each axis, coloured by valence.

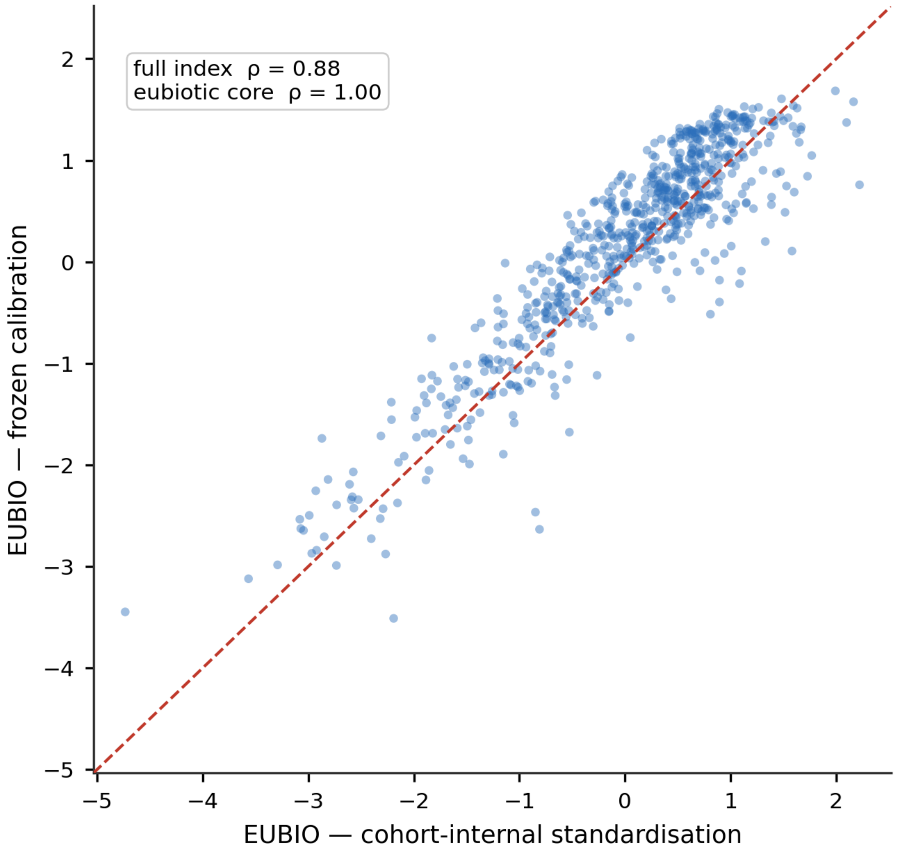


**Extended Data Fig. 3 | The ranking is calibration-independent..** EUBIO computed with the frozen calibration against a cohort-internal standardisation on the public PD cohort; the eubiotic core is perfectly scale-invariant ($\rho$ = 1.00) and the full index nearly so ($\rho$ = 0.88), so no reported ranking depends on the proprietary reference.

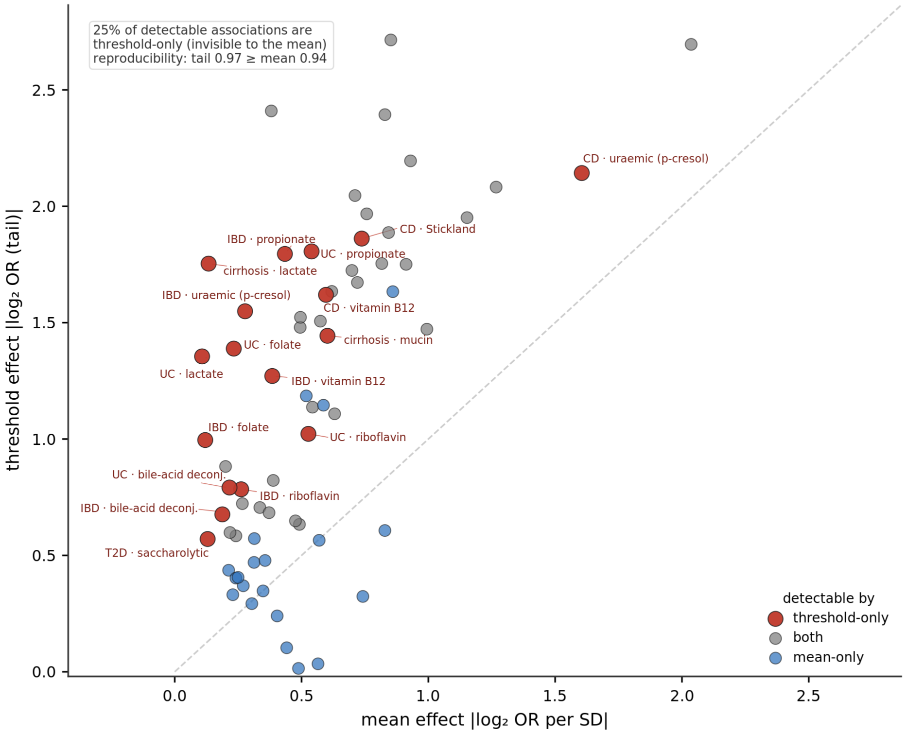


**Extended Data Fig. 4 | Dysbiosis is often a threshold phenomenon..** Each detectable channel-by-condition association in cMD, plotted by the magnitude of its mean effect (odds ratio per standard deviation) against its threshold effect (odds ratio for occupying the dysbiotic tail). Associations detectable only through the threshold (red) sit far from the mean axis; about a quarter of detectable associations are of this kind, and they reproduce across studies at least as well as mean associations (sign concordance 0.97 against 0.94).

### Supplementary Tables

#### Table S1 | Causal-anchor validation.

Per-cohort EUBIO response under antibiotics and faecal microbiota transplantation, with the reproducibility outcome.

| Figure | Metric | Cohort | EUBIO | Outcome |
|---|---|---|---|---|
| Fig2 | paired Δ EUBIO | PRJEB20800 | -4.13 (12/12, n=12) | drop |
| Fig2 | paired Δ EUBIO | PRJEB28058 | -0.68 (31/41, n=41) | drop |
| Fig2 | paired Δ EUBIO | PRJEB47909 | +3.23 (11/11, n=11) | recovery |
| Fig2 | paired Δ EUBIO | PRJNA525458 | +2.85 (6/8, n=8) | recovery |
| Fig2 | paired Δ EUBIO | PRJNA701961 | +3.54 (6/7, n=7) | recovery |
| Fig2 | paired Δ EUBIO | PRJEB39023 | +2.47 (7/8, n=8) | recovery |
| Fig2 | paired Δ EUBIO | PRJNA628604 | +0.40 (9/13, n=13) | recovery |
| Fig2 | paired Δ EUBIO | PRJEB44237 | -0.01 (11/24, n=24) | changed |
| Fig2 | paired Δ EUBIO | PRJNA285502 | -0.01 (1/6, n=6) | drop |
| Fig3B | specificity resp-vs-non (MWU p) | PRJNA672867 | +0.41 vs +0.46, p=0.84 | correctly non-separating |
| Fig3B | donor eubiosis (mean EUBIO) | PRJNA672867 | +0.41 | donors mildly eubiotic |
| Fig4A | AUC eubiotic vs dysbiotic | pooled anchors | 0.668 | changed |
| Fig4A | AUC eubiotic vs dysbiotic | pooled anchors | 0.680 | diversity (context) |
| Fig4C | AUC Koopen | PRJEB44237 | 0.637 | EUBIO discriminates where Shannon fails |
| Fig4C | AUC Koopen | PRJEB44237 | 0.492 | Shannon ~chance |
| SuppS2 | within-subj dz (baseline→nadir) | PRJEB20800 | EUBIO dz=-2.77 | abx drops eubiosis |
| SuppS2 | within-subj dz (baseline→nadir) | PRJEB28058 | EUBIO dz=-0.25 | abx drops eubiosis |
| Fig3A | carbapenemase EC3.5.2.6 resp vs non (MWU p) | PRJNA628604 | P = 0.011 | exact match |
| Fig3C | reliability ICC(1) | PRJNA664754 | 0.47 (gated, >500 EC) | reproduced (ICC 0.47) |

#### Table S2 | Channel-by-condition effect sizes.

Covariate-adjusted, within-study Hedges' $g$ for each functional channel across six conditions of cMD (positive = higher in cases), with channel valence. These are continuous (mean) effect sizes and are small by design: because most of the disease signal is a threshold rather than a shift in the mean, the corresponding gate odds ratios are typically much larger (Table S3; Extended Data Fig. 4).

| Channel | CRC | IBD | CD | UC | T2D | cirrhosis | valence |
|---|---|---|---|---|---|---|---|
| Redox engine | 0.05 | 0.013 | 0.006 | 0.037 | 0.073 | 0.043 | protective |
| Tryptophan-indole | 0.059 | 0.022 | 0.013 | 0.022 | 0.03 | 0.075 | context |
| Polyamines | 0.015 | 0.053 | 0.067 | 0.059 | -0.009 | 0.08 | context |
| Histamine | 0.035 | 0.027 | 0.067 | 0.025 | 0.002 | 0.047 | context |
| Propionate (succinate) | 0.124 | 0.002 | -0.057 | 0.019 | -0.001 | 0.03 | context |
| Lactate | -0.054 | 0.017 | -0.011 | 0.051 | 0.018 | 0.052 | danger |
| Bile-acid deconjugation | -0.037 | 0.038 | 0.073 | 0.034 | 0.032 | -0.035 | context |
| Secondary bile acid | 0.031 | 0.022 | 0.074 | 0.014 | 0.063 | 0.061 | context |
| Trimethylamine | 0.063 | 0.073 | 0.098 | 0.09 | 0.029 | 0.121 | danger |
| Uraemic (p-cresol) | 0.005 | 0.049 | 0.128 | 0.088 | 0.066 | 0.069 | danger |
| Stickland proteolysis | 0.044 | 0.058 | 0.135 | 0.029 | 0.045 | 0.007 | danger |

| Channel | CRC | IBD | CD | UC | T2D | cirrhosis | valence |
|---|---|---|---|---|---|---|---|
| Hydrogen sulfide | 0.06 | -0.012 | 0.03 | -0.019 | 0.053 | 0.003 | context |
| Vitamin B12 | -0.023 | 0.002 | 0.067 | -0.012 | -0.013 | 0.03 | context |
| Vitamin K2 | 0.066 | 0.045 | 0.098 | -0.005 | -0.024 | 0.134 | context |
| Riboflavin | -0.003 | -0.045 | -0.005 | -0.092 | 0.013 | 0.068 | protective |
| Folate | -0.03 | 0.028 | 0.03 | 0.024 | 0.055 | 0.114 | context |
| Mucin foraging | 0.024 | 0.016 | 0.045 | -0.006 | 0.024 | 0.103 | context |
| Ethanolamine | -0 | 0.069 | 0.154 | 0.132 | 0.03 | 0.101 | danger |
| LPS endotoxin | 0.007 | 0.025 | 0.066 | 0.03 | 0.017 | 0.127 | danger |
| Saccharolytic | -0.051 | -0.021 | -0.021 | -0.025 | -0.042 | 0.019 | protective |
| Glycine cleavage | 0.615 | -0.092 | 0.233 | -0.242 | -0.114 | 0.261 | context |

#### Table S3 | The engine as a threshold: continuous versus gate.

For each condition, the engine's sink-to-respiration log-ratio read continuously (odds ratio per standard deviation) and as a threshold (odds ratio for occupying the oxidised niche), with the fraction of cases and controls in that niche. Tail phenomena (continuous null, gate significant) are the rule for inflammatory and hepatic disease.

| Condition | continuous OR/SD | cont. P | gate OR | gate P | % cases oxidised | % controls oxidised | n studies | evidence |
|---|---|---|---|---|---|---|---|---|
| CRC | 1.047 | 0.506 | 1.084 | 0.255 | 18.6 | 13.4 | 11 | strong |
| adenoma | 0.938 | 0.62 | 1.187 | 0.111 | 13.8 | 6 | 5 | strong |
| IBD | 0.796 | 0.143 | 2.034 | 0 | 10.6 | 3.9 | 3 | strong |
| UC | 0.873 | 0.41 | 2.332 | 0.052 | 11.6 | 3.9 | 3 | strong |
| CD | 0.749 | 0.369 | 1.589 | 0.076 | 10.3 | 3.9 | 3 | strong |
| cirrhosis | 0.413 | 0 | 1.848 | 0 | 48 | 16.6 | 2 | association |
| STH | 0.479 | 0.001 | 2.01 | 0.001 | 48.1 | 23.7 | 1 | weak |
| T2D | 0.936 | 0.57 | 1.016 | 0.877 | 12.2 | 7.7 | 5 | strong |
| IGT | 0.932 | 0.264 | 1.143 | 0.047 | 5.2 | 4.1 | 3 | strong |
| ACVD | 1.278 | 0.116 | 0.938 | 0.625 | 27.6 | 22.8 | 1 | weak |
| hypertension | 0.71 | 0.113 | 1.325 | 0.163 | 59.6 | 51.2 | 1 | weak |
| schizophrenia | 0.884 | 0.481 | 0.979 | 0.901 | 35.5 | 33.3 | 1 | weak |
| ME/CFS | 0.802 | 0.366 | 0.943 | 0.807 | 12 | 10 | 1 | weak |

#### Table S4 | Cross-population EUBIO, diversity and a beneficial-taxa index.

EUBIO (median and interquartile range) with Shannon diversity and a fixed-panel beneficial-taxa index for the same populations; diversity and the taxa panel do not track the functional ranking.

| Population | n (usable) | median EC detected | EUBIO median | IQR | Shannon | beneficial-taxa index | Coverage note |
|---|---|---|---|---|---|---|---|
| Japan (contemporary healthy) | 384 | 619 | 0.43 | [-0.14,0.88] | 2.98 | 34.5 | - |
| Hadza (Tanzania hunter-gatherer) | 40 | 437 | 1.56 | [0.12,2.02] | 1.56 | 6.1 | - |
| Sardinia young (21-33) | 17 | 579 | 1.11 | [0.32,1.36] | 3.31 | 44.3 | - |

| Population | n (usable) | median EC detected | EUBIO median | IQR | Shannon | beneficial-taxa index | Coverage note |
|---|---|---|---|---|---|---|---|
| Sardinia elderly (68-88) | 23 | 603 | 0.56 | [-0.34,1.05] | 3.27 | 35.2 | - |
| Sardinia centenarians (99-107) | 19 | 1135 | -0.92 | [-1.32,-0.00] | 3.5 | 36.2 | - |
| Tunapuco+Matsés (Peru traditional) | 13 | 460 | 1.51 | [1.15,1.70] | 2.98 | 19.2 | 13/72 samples above coverage floor |

#### Table S5 | TAGMOS architecture composition.

Each axis with its scoring type, valence, reliability tier (A/B/C, where assigned) and the number of annotation-verified enzymes it aggregates. Enzyme identities are proprietary and are not released.

| Axis | axis type | valence | tier | n enzymes |
|---|---|---|---|---|
| Redox engine | balance | protective | A | 11 |
| Riboflavin | abundance | protective | A | 4 |
| Saccharolytic | abundance | protective | A | 4 |
| Mucin foraging | abundance | context | - | 4 |
| Hydrogen sulfide | balance | context | - | 3 |
| Polyamines | abundance | context | C | 3 |
| Secondary bile acid | abundance | context | B | 2 |
| Folate | abundance | context | C | 2 |
| Glycine cleavage | abundance | context | A | 2 |
| Propionate | abundance | context | B | 2 |
| Vitamin B12 | abundance | context | B | 2 |
| Vitamin K2 | abundance | context | B | 2 |
| Bile-acid deconjugation | abundance | context | B | 1 |
| Histamine | presence flag | context | C | 1 |
| Tryptophan-indole | abundance | context | - | 1 |
| Trimethylamine | abundance | danger | A | 3 |
| D-lactate | balance | danger | A | 2 |
| LPS endotoxin | abundance | danger | A | 2 |
| Stickland proteolysis | abundance | danger | A | 2 |
| Ethanolamine | abundance | danger | C | 1 |
| Uraemic (p-cresol) | abundance | danger | C | 1 |

#### Table S6 | Threshold versus mean detection is robust.

Across gate thresholds and false-discovery cutoffs, the number of channel-by-condition associations detectable by the mean, the threshold, or both; the proportion detectable only through the threshold is stable at roughly a quarter to a third, and the cross-study sign concordance of the threshold reading matches or exceeds that of the mean.

| gate threshold (SD) | FDR q | detectable | both | mean-only | threshold-only | % threshold-only | concord. threshold | concord. mean |
|---|---|---|---|---|---|---|---|---|
| 1 | 0.05 | 51 | 20 | 16 | 15 | 29.4 | 0.99 | 0.96 |
| 1 | 0.1 | 67 | 30 | 20 | 17 | 25.4 | 0.97 | 0.94 |

| gate threshold (SD) | FDR q | detectable | both | mean-only | threshold-only | % threshold-only | concord. threshold | concord. mean |
|---|---|---|---|---|---|---|---|---|
| 1.5 | 0.05 | 59 | 19 | 17 | 23 | 39 | 0.87 | 0.96 |
| 1.5 | 0.1 | 71 | 25 | 25 | 21 | 29.6 | 0.88 | 0.94 |
| 2 | 0.05 | 45 | 11 | 25 | 9 | 20 | 0.87 | 0.96 |
| 2 | 0.1 | 57 | 14 | 36 | 7 | 12.3 | 0.88 | 0.94 |
